# Mapping the T cell repertoire to a complex gut bacterial community

**DOI:** 10.1101/2022.05.04.490632

**Authors:** Kazuki Nagashima, Aishan Zhao, Katayoon Atabakhsh, Allison Weakley, Sunit Jain, Xiandong Meng, Alice G. Cheng, Min Wang, Steven Higginbottom, Alex Dimas, Pallavi Murugkar, Michael A. Fischbach

## Abstract

Certain bacterial strains from the microbiome induce a potent, antigen-specific T cell response^1–5^. However, the specificity of microbiome-induced T cells has not been explored at the strain level across the gut community. Here, we colonize germ-free mice with a complex defined community (97 or 112 bacterial strains) and profile T cell responses to each strain individually. Unexpectedly, the pattern of T cell responses suggests that many T cells in the gut repertoire recognize multiple bacterial strains from the community. We constructed T cell hybridomas from 92 T cell receptor (TCR) clonotypes; by screening every strain in the community against each hybridoma, we find that nearly all of the bacteria-specific TCRs exhibit a one-to-many TCR-to-strain relationship, including 13 abundant TCR clonotypes that are polyspecific for 18 Firmicutes in the community. By screening three pooled bacterial genomic libraries against 13 pooled hybridomas, we discover that they share a single target: a conserved substrate-binding protein (SBP) from an ABC transport system. Treg and Th17 cells specific for an epitope from this protein are abundant in community-colonized and specific-pathogen-free mice. Our work reveals that T cell recognition of Firmicutes is focused on a widely conserved cell-surface antigen, opening the door to new therapeutic strategies in which colonist-specific immune responses are rationally altered or redirected.

## INTRODUCTION

Immune modulation by the gut microbiome plays an important role in a variety of diseases and therapeutic indications^5–7^. As a result, strategies for controlling or redirecting colonist-specific immune responses have considerable therapeutic promise. However, progress toward this goal is impeded by an incomplete understanding of the logic underlying immune recognition of the microbiome.

In pioneering efforts to date, microbiome immunology has been characterized at two levels: the community and the strain. Transplanting human fecal communities into germ-free mice impacts a variety of immune-linked disease phenotypes including inflammatory bowel disease, obesity, autism, malnutrition, and the response to cancer immunotherapy^8–14^, but identifying the causative strains has been challenging.

At the strain level, two themes have emerged. First, the nature of the strain determines what type of immune cell is induced; for example, the gut colonists segmented filamentous bacterium (SFB)^15^ and *Helicobacter*^16,17^ induce Th17 cells and regulatory T cells, respectively. Second, in many cases, the immune cells elicited have a T or B cell receptor specific for an epitope from the strain that elicited them^1–4,18,19^. These studies have led to the hypothesis that a few “keystone” species dominantly influence the repertoire and function of gut T cells^5,20^.

Efforts have been made to bridge community- and strain-level analysis. In a series of pioneering studies, strains responsible for immune modulation have been identified from undefined communities^21–24^; in another effort, a large set of isolates were profiled under conditions of mono-colonization^25^. Both approaches identify strains that are capable of modulating immune function, but neither one reveals how a strain behaves in the context of a complex community. Notably, the phenotype of T cells induced by *Akkermansia muciniphila* changes depending on the complexity of the community in which it resides^4^; it is unclear whether the same is true for other common species. Moreover, it is unknown what each strain in the community contributes to the ‘sum total’ phenotype of immune modulation by the microbiome.

Here, we address these questions by introducing a new system for studying interactions between the immune system and a microbiome that is defined, but complex enough to be physiologically relevant. We colonize germ-free mice with this community (97 or 112 bacterial strains) and profile T cell responses to each strain individually. We found that the pattern of T cell responses is not consistent with a model in which each bacterial strain stimulates its own antigen-specific T cell population. Using a combination of single-cell RNA and TCR sequencing (scRNA-seq and scTCR-seq), we identified microbiome-responsive TCR clonotypes. By generating 92 T cell hybridomas and screening each one against every strain in the community, we show that expanded, microbiome-responsive TCR clonotypes are often specific for multiple strains in the community—i.e., there is a one-to-many TCR-to-strain mapping. 13 such clonotypes recognize 18 Firmicutes from the community; by screening genomic libraries from three of the Firmicutes, we show that they recognize a single antigen: a substrate-binding protein (SBP) conserved among Firmicutes that functions as part of an ABC transport system. Our results suggest that immune surveillance of the gut microbiome relies on expanded T cell clonotypes that play ‘zone defense’ by recognizing extracellular antigens conserved among widely divergent species.

### Complex defined community as a model for immune modulation by the gut microbiome

As a starting point for our work, we set out to determine whether germ-free mice colonized by a complex defined community have a similar profile of T cell subtypes to that of conventionally colonized mice. To address this question, we colonized germ-free C57BL/6 mice with a 97- or 112-member gut bacterial community (hereafter, hCom1d and hCom2d) (**Supplementary Table 1**), derivatives of gut bacterial communities recently developed as a defined model system for the gut microbiome^26^. After two weeks of colonization, we isolated intestinal T cells and profiled them by flow cytometry (**Fig. 1a**). We find that the T cell profile of hCom1d- and hCom2d-colonized mice is similar to that of conventionally colonized (specific pathogen free, SPF) mice and distinct from that of germ-free mice (**Fig. 1b**). We conclude that hCom1d- and hCom2d-colonized mice are a reasonable setting in which to study immune modulation by the gut microbiome.

**Figure 1:**
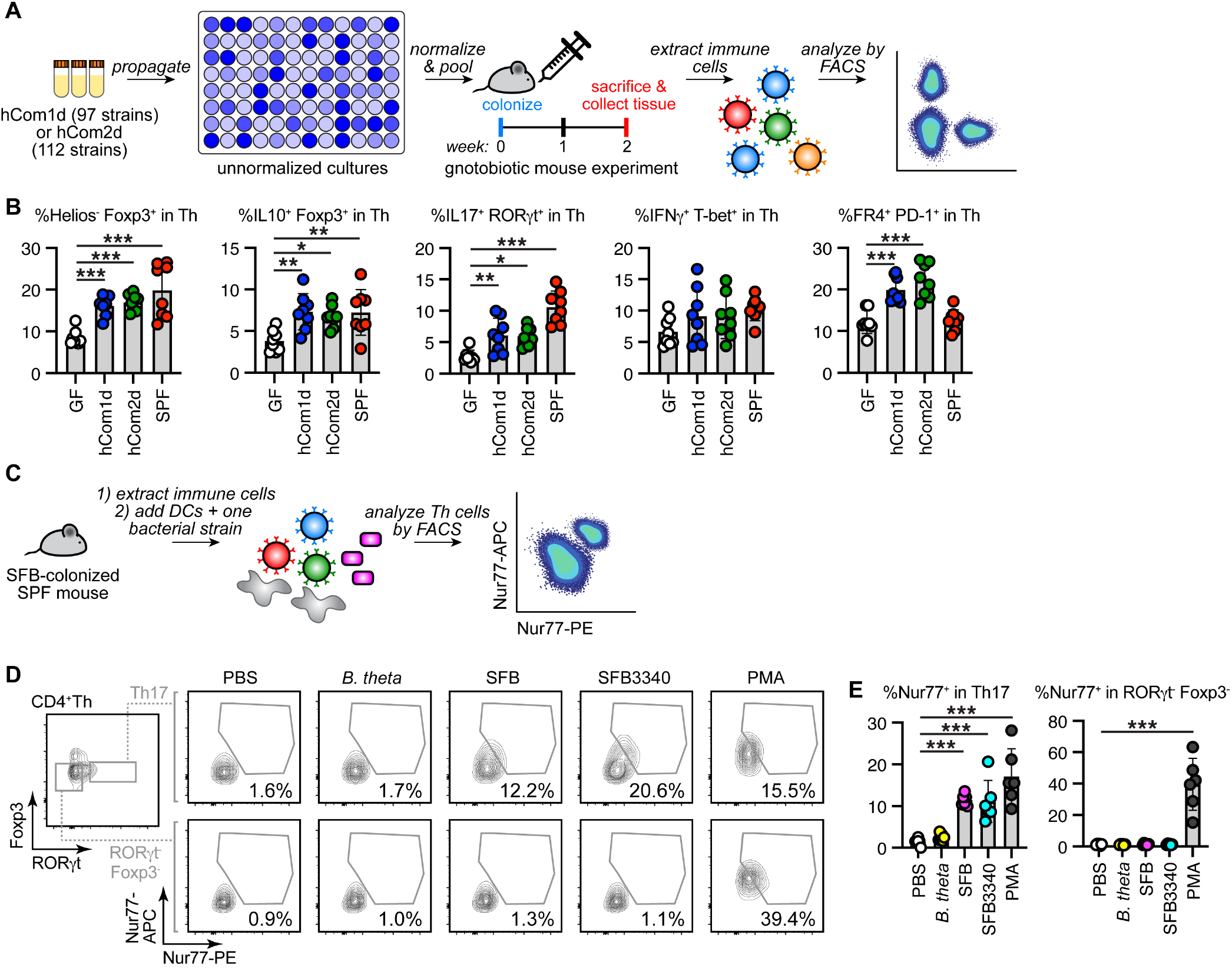
A model system for studying immune modulation by the gut microbiome. (**A**) Schematic of the T cell profiling experiment. Frozen stocks of 97 strains (hCom1d) or 112 strains (hCom2d) were used to inoculate cultures that were grown for 48 h, diluted to similar optical densities, and pooled. The mixed culture was used to colonize germ-free C57BL/6 mice by oral gavage. Mice were housed for two weeks before sacrifice. Immune cells from the large intestine were extracted, stimulated by PMA/ionomycin and analyzed by flow cytometry. (**B**) Th cell subtypes, as a percentage of the total Th cell pool, were broadly similar among hCom1d-colonized, hCom2d-colonized, and SPF mice and distinct from germ-free mice. *n*=8 mice per group. (**C**) Schematic of the mixed lymphocyte assay. SFB-colonized SPF mice were sacrificed; immune cells from the large intestine were extracted and co-cultured with a heat-treated bacterial strain and dendritic cells.After 4 h, cells were fixed, stained with two antibodies specific for Nur77 and analyzed by flow cytometry. (**D**) Gating strategy for the mixed lymphocyte assay. Expression of Nur77 was analyzed in Th17 cells and RORgt^-^ Foxp3^-^ Th cells to evaluate TCR stimulation. Th17 cells were stimulated by SFB (fecal pellets from SFB-mono-colonized mice), purified SFB3340 peptide (an antigen from SFB) and PMA/ionomycin, a positive control. (**E**) Statistical analysis for the mixed lymphocyte assay. *n*=6 mice per group. Statistical significance was assessed using a one-way ANOVA in (**B**) and (**E**) (\**p*<0.05; \*\**p*<0.01; \*\*\**p*<0.001). Data shown are mean ± standard deviations from 2 independent experiments in (**B**) and (**E**).

### Profiling T cell responses to each strain in the community

Next, we sought to understand how each strain modulates T cell immunity in the physiologic setting of a native-scale community. We envisioned an experiment in which T cells isolated from hCom1d-colonized mice could be incubated in vitro with each bacterial strain in the community—one at a time— plus dendritic cells for antigen presentation. Since primary T cells can change state while being cultured ex vivo for an extended period of time, a requirement for this experiment is an assay that would give us a rapid, sensitive readout of primary T cell stimulation. To this end, we established a new assay in which we measure the level of Nur77, an early marker of TCR stimulation^27,28^. We found that monitoring Nur77 expression with two distinct antibodies constituted a sensitive and specific assay for primary T cell stimulation (**Fig. 1c**). We calibrated this assay using a well-established commensal-specific T cell response, Th17 cell induction by SFB^1^. T helper (Th) cells were extracted from Taconic mice harboring SFB and co-cultured with antigens and antigen-presenting cells (**Fig. 1d,e**). We observed the upregulation of Nur77 only in Th17 cells co-cultured with SFB or the SFB antigen SFB3340, whereas PMA stimulated Nur77 expression to a similar extent in Th17 cells and ROR*γ*t^-^ Th cells, demonstrating that the co-culture assay is sufficiently sensitive and specific to identify interactions between commensal bacterial strains and antigen-specific T cells.

To determine which T cell subtypes are restimulated by strains in the community, we colonized germ-free C57BL/6 mice with hCom1d, waited two weeks to give naïve T cells time to differentiate, and then isolated T cells from the large intestine. We incubated these T cells individually with each of the 97 strains in hCom1d, using murine dendritic cells for antigen presentation (**Fig. 2a**); as a control, we used germ-free mice from which T cells were harvested and profiled in the same manner. We measured the proportion of regulatory T (Treg) cells, Th17 cells, and FR4^+^ Th cells restimulated by each strain using flow cytometry. Most of the strains in the community did not restimulate T cells from germ-free mice (**Fig. 2b**). In contrast, the majority of the strains in the community (70/97 strains, p<0.05) restimulated at least one type of T cell from hCom1d-colonized mice, and more than a third of the strains restimulated multiple T cell subtypes (31/97). More strains activate Tregs (51) than Th17 (36) or FR4^+^ Th (21) cells. *A. muciniphila* is known to induce a context-dependent T cell response: predominantly T follicular helper cells in the presence of a simple defined community, and other effector Th cells in the presence of a more complex SPF microbiome^4^. We observed that the intestinal T cell pool in hCom1d-colonized mice contains *A. muciniphila*-specific Treg and Th17 cells, providing further evidence that hCom1d mimics the function of the complex gut microbiome.

**Figure 2:**
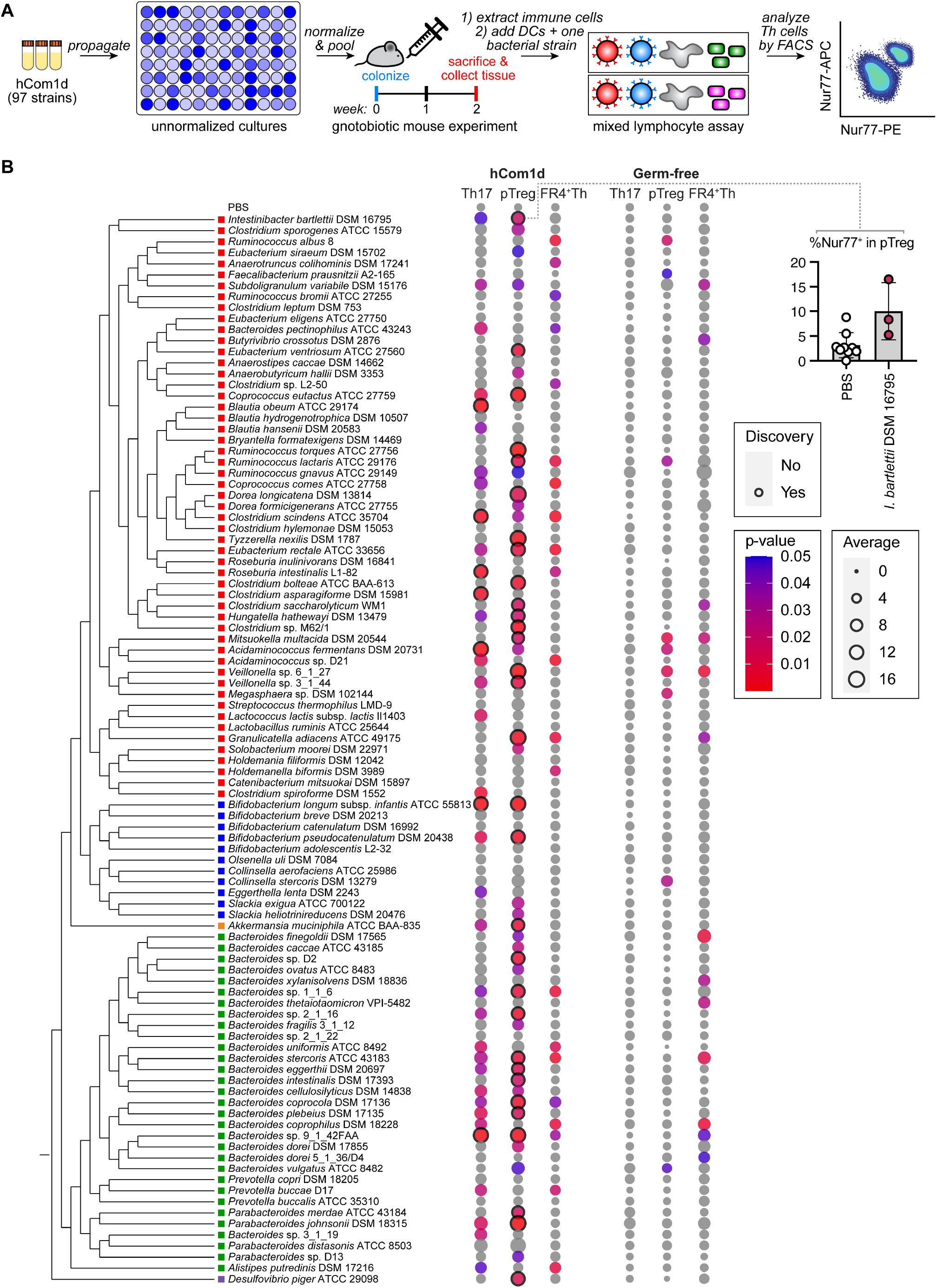
Strain-by-strain profiling of T cell responses to a complex defined community. (**A**) Schematic of the experiment. Frozen stocks of the 97 strains from hCom1d were used to inoculate cultures that were grown for 48 h, diluted to similar optical densities, and pooled. The mixed culture was used to colonize germ-free C57BL/6 mice by oral gavage. After two weeks, 10-15 mice were sacrificed to isolate T cells from the large intestine. T cells from the mice were pooled and co-cultured with each of the 97 heat-treated strains in hCom1d, along with dendritic cells for antigen presentation. After 4 hours, T cells were fixed and stained for Nur77 expression and analyzed by flow cytometry. As a negative control, T cells from germ-free mice were co-cultured and profiled in the same manner. (**B**) Profiling T cell reactivity against each of hCom1d strains. Dot size shows the average percentage of Nur77^+^ cells in a T cell subset after coculture. Dot color represents p-value in comparison with the negative control, treatment with PBS. Data showing the restimulation of pTreg cells by *Intestinibacter bartlettii* DSM 16795 are displayed as an example (upper right). A phylogenetic tree of the strains in hCom1d was generated based on a multiple sequence alignment generated from conserved single-copy genes. The colored square to the left of each strain name indicates its phylum: Firmicutes = red, Actinobacteria = blue, Verrucomicrobia = orange, Bacteroidetes = green, and Proteobacteria = purple. *n*=9 replicates were used for PBS and *n*=3 replicates were used for the remaining samples. The data represent 3 independent experiments per colonization condition. For statistical analysis, multiple comparison testing was performed by GraphPad Prism using the two-stage linear step-up method (Benjamini, Krieger and Yekutieli) for controlling the false discovery rate.

If each strain induced its own ‘private’ T cell subpopulation, the sum over all the T cells stimulated by each strain would be ≤100% of the gut T cell population. Intriguingly, the sum over all the stimulated T cells is far greater than 100% (**Fig. 2b**): Nur77^+^ pTreg, Th17, and FR4^+^ Th cells = 863%, 811%, and 536%, respectively. We hypothesized that this pattern of restimulation arises from a scenario in which T cells are ‘polyspecific’ to multiple strains, and the proportion of cells exceeding the 100% threshold is a consequence of multiple strains restimulating the same T cell clonotypes.

### Identifying microbiome-responsive T cell clonotypes

To test this hypothesis, we sought to identify T cell clonotypes—individual cells or groups of cells that express the same TCR—that are responsive to the gut microbiome so we could generate a higher-resolution map of strains to clonotypes. We colonized germ-free mice with hCom1d or hCom2d. We pooled three mice per condition, isolated immune cells from the small and large intestine, enriched the CD3^+^fraction by fluorescence-activated cell sorting (FACS), and analyzed these cells using a combination of scRNA-seq and scTCR-seq (**Fig. 3a**). The dataset of transcriptomes from 35,237 cells (4,908-7,029 cells per condition) were filtered, normalized and analyzed by uniform manifold approximation and projection (UMAP) (**Fig. 3b**). Unbiased clustering defined 25 clusters (**Extended Data Fig. 1a**). We visualized the expression of canonical subset markers and matched the 25 clusters to 16 cell types (**Extended Data Fig. 1b-d** and **Fig. 3b**). T effector cell subsets showed a continuous distribution, consistent with a recent report^29^. The FR4^+^ Th cell type, which corresponds to FR4^+^ PD-1^+^ Th cells in **Fig. 1a**, expresses Il21 and Cxcr5 and may affect B cell class switching (**Extended Data Fig. 1b-d**). The ‘other effector Th’ cell type does not express Th-subset-defining markers but exhibited high expression of Il21r, Bcl2a1 and Tgfb1, suggesting that this uncharacterized population may play a distinct role in intestinal immunity (**Extended Data Fig. 1b-d**). We found that colonization by hCom1d or hCom2d changed the composition of immune cell subsets more profoundly in the large intestine than the small intestine; colonic pTreg, Th17 and FR4^+^ Th were induced (**Extended Data Fig. 2a, b**), consistent with our initial findings (**Fig. 1b**).

**Figure 3:**
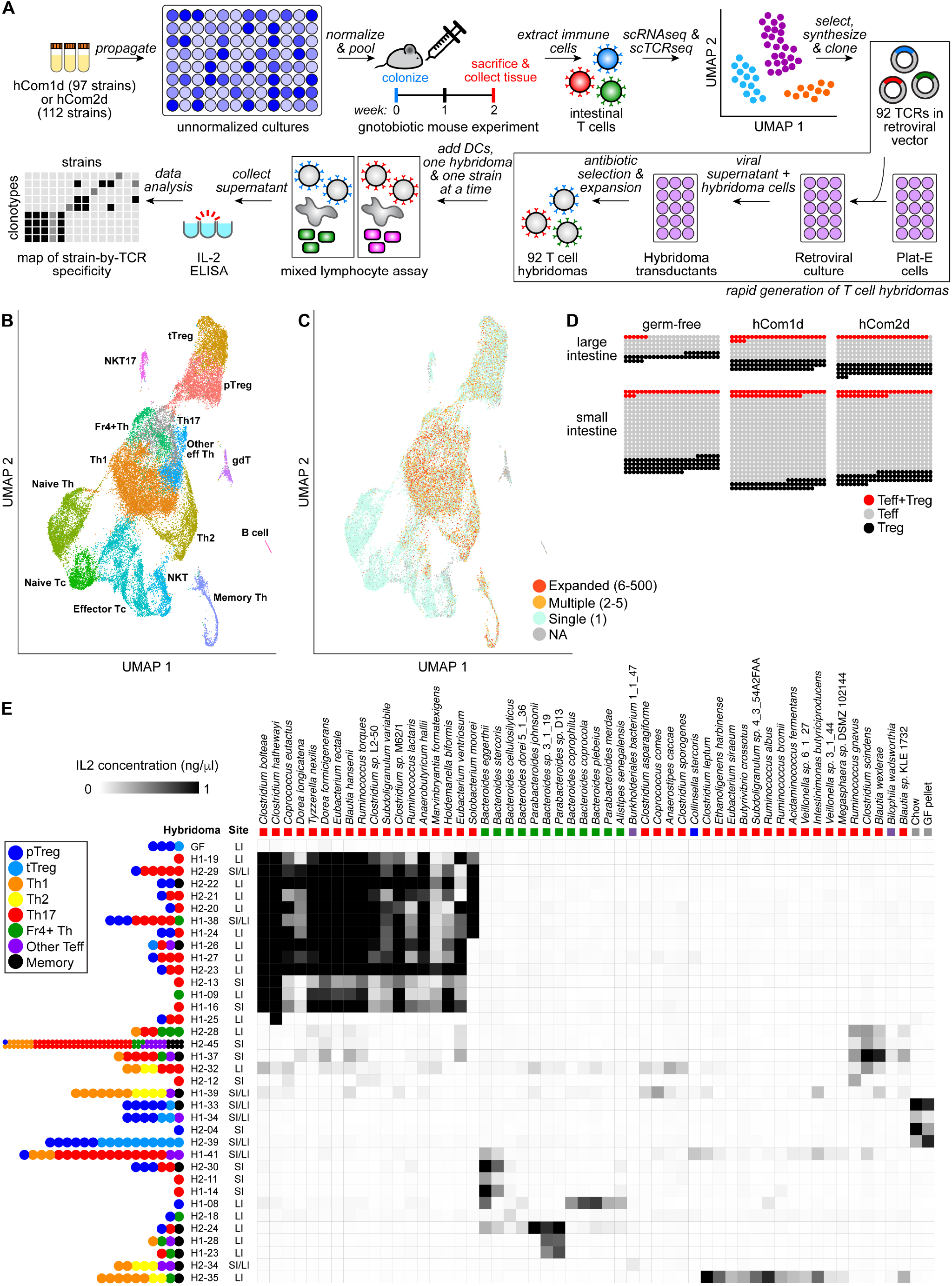
scRNA-seq and scTCR-seq to identify microbiome-responsive T cell clonotypes. (**A**) Schematic of the experiment. Frozen stocks of the 97 strains from hCom1d were used to inoculate cultures that were grown for 48 h, diluted to similar optical densities, and pooled. The mixed culture was used to colonize germ-free C57BL/6 mice by oral gavage. After two weeks, intestinal T cells were isolated, purified, and analyzed by scRNA-seq and scTCR-seq. For generating TCR hybridomas, 92 TCR clonotypes were selected based on one of two criteria: (*i*) we chose 55 clonotypes that were ‘expanded’ in that they occurred >2 times in our combined pool of 35,237 cells (range=2-84). (*ii*) We selected 37 clonotypes that harbored an expression signature consistent with being microbiome-specific (see **Extended Data Fig. 2c, d**). Synthetic expression constructs that consist of the TCR a and β chains separated by a self-cleaving P2A peptide were cloned into the retroviral vector pMSCV-mCD4-PIG and used to transfect Plat-E cells. Retrovirus was collected from these cells and used to transduce NFAT-GFP hybridoma cells, which were enriched for transductants using a selectable marker. Each T cell hybridoma was co-cultured with every strain in hCom1d and hCom2d, one strain at a time. We measured IL-2 production by the T cell hybridomas to detect TCR stimulation. Data from this experiment were analyzed to create a map of strain-TCR specificity. (**B**) Uniform manifold approximation and projection (UMAP) plot. These data represent three merged samples: hCom1d-colonized, hCom2d-colonized, and germ-free mice. See **Extended Data Fig. 1** for gene expression profiling to assign clusters to T cell subsets. (**C**) Frequency of TCR clonotypes on the UMAP plot. Expanded TCRs (red) represent clonotypes observed in more than 5 cells; multiple (yellow) are clonotypes found in 2-5 cells; and single (light blue) were seen in only one cell. Most of the expanded TCR clonotypes have an expression profile consistent with T effector cells, whereas naïve T cells are rich in unique (i.e., non-expanded) TCR clonotypes. (**D**) Analysis of expanded TCR clonotypes in each sample. Each dot represents one TCR clonotype found in multiple T cells (red = shared between T effector cells and Treg cells; gray = T effector; black = Treg). (**E**) A map of TCR-strain specificity. Each T cell hybridoma was co-cultured with every strain individually. Shown are the 35 TCRs reactive to at least one bacterial strain and the 55 antigens (53 strains, chow, and a GF fecal pellet) that stimulated at least one TCR. 13 T cell hybridomas were reactive to a subset of 15-18 Firmicutes. GF, a T cell hybridoma derived from germ-free mice, is shown as a negative control. Colored dots at left represent the primary T cells in which the corresponding TCR was expressed. H1-## indicates hybridomas expressing a TCR selected from hCom1d-colonized mice, and H2-## are TCR hybridomas from hCom2d-colonized mice. The colored square below each strain name indicates its phylum: Firmicutes = red, Bacteroidetes = green, and Proteobacteria = purple. SI = small intestine, LI = large intestine. The data are an average of two independent experiments.

To explore the pattern of TCR specificities across T cell subsets, we combined the scRNA-seq and scTCR-seq data. The pattern of TCR clonotypes was consistent with a well-established model: after a naïve T cell differentiates into a T effector cell or a Treg, it clonally expands (**Fig. 3c**). Notably, the frequency of clonotypes in which the same TCR is found on a Treg and a T effector cell increases when the mice are colonized (**Fig. 3d**).

With the combined scRNA-seq and scTCR-seq data in hand, we sought to determine how microbiome-specific TCRs map to strains from the community. We reasoned that if we could identify TCR clonotypes from the broader pool that were specific for the microbiome, the corresponding TCR genes could be used to construct T cell hybridomas, enabling us to assay TCR specificity against each strain from the community (**Fig. 3a**). We used two orthogonal criteria to select a group of TCR clonotypes that were likely to be microbiome-specific (**Supplementary Table 2**):

First, we chose 55 clonotypes that were ‘expanded’ in that they occurred >2 times in our combined pool of 35,237 cells (range=2-84). We reasoned that expanded TCR clonotypes may have derived from T cell clones that divided in response to bacterial stimulation. Moreover, they may be functionally important given their prevalence in the repertoire. Clonotypes that represent a TCR shared by Treg and T effector cells were of particular interest since they increase after colonization.

Second, we analyzed scRNAseq data to identify genes that were differentially expressed in community-colonized versus germ-free mice (**Extended Data Fig. 2c**). We used these colonization-induced or repressed genes together with T cell subset markers to choose 37 clonotypes that harbored an expression signature consistent with being microbiome-specific (**Extended Data Fig. 2d** and **Supplementary Table 2**). For both categories, clonotypes were selected to represent a mixture of tissue origins (small vs. large intestine) and T cell subtypes (Treg, FR4^+^ Th, Th17, Th1, Th2 or a combination thereof).

### Constructing and assaying hybridomas to determine how strains map to TCR clonotypes

To determine how strains map to TCR clonotypes, we constructed 92 T cell hybridomas, each one expressing a single TCR. To construct hybridomas en masse (**Fig. 3a**), we ordered synthetic expression constructs that consist of the TCR a and β chains separated by a self-cleaving P2A peptide. These constructs were cloned into a retroviral vector and propagated in Plat-E cells in a 96-well format; the virus-containing supernatant was used to transduce the NFAT-GFP hybridoma cell line^30^. We used puromycin to enrich for transductants and used the enriched pools for the experiments described below.

A variety of T cell epitopes from other gut commensals have been discovered and characterized^1,2,4,31–35^. We started by testing 20 such antigens against the 92 hybridomas using dendritic cells for antigen presentation (**Extended Data Fig. 3**). After overnight coculture, we measured the concentration of IL-2 in the culture supernatant as a reporter of TCR stimulation and signal transduction. None of the hybridomas reacted to any of the antigens, indicating that the hybridomas are not specific for a previously discovered epitope.

Next, we incubated each hybridoma with every strain individually (120 antigens x 92 hybridomas = 11,040 assays) (**Fig. 3e**). ~30% of the T cell hybridomas (31/92) were responsive to at least one strain, and ~45% of the bacterial strains (53/118) stimulated at least one TCR. Four additional hybridomas were stimulated by homogenized mouse chow, indicating that they are food responsive; the remainder are not responsive to the microbiome or food. We confirmed this result by an independent repeat of the co-culture assays using the microbiota- and food-reactive hybridomas (55 antigens x 36 hybridomas = 1,980 assays). TCR sequences selected from the ‘expanded’ clonotypes were more responsive to bacterial strains or food (26/55 hybridomas) than clonotypes selected for an apparent microbiome-induced gene signature (9/37 hybridomas).

The hybridoma-strain specificity data reveals clusters of 3-5 TCRs that are specific for 2-4 strains in the community (or for chow). There are apparent similarities among the cell type(s) of origin of the TCRs, and phylogenetic similarities among the strains for which they are specific. These data represent only a miniscule portion of the T cell repertoire, so no broad conclusions about frequency or absolute numbers can be drawn from this experiment. Nonetheless, it is striking that even in a portion of the repertoire this small, TCRs appear to be generally polyspecific to multiple bacterial strains.

In addition to these smaller clusters, we were intrigued by the observation that 13 of the microbiome-responsive hybridomas were stimulated by a subset of 15-18 Firmicutes. These are not the most abundant strains in hCom1d-colonized mice (**Extended Data Fig. 4**), so this is not a trivial consequence of bacterial cell numbers in vivo. Many of these clonotypes were found simultaneously in Treg and effector T cells in the population analyzed by scRNAseq, indicating that simultaneous restimulation of Treg/Teff observed in the mixed lymphocyte screen occurs not justat the population level but within individual T cell clonotypes. These data are consistent with the possibility that there exist abundant, polyspecific T cells that target multiple strains of Firmicutes.

### Identifying the antigen targeted by the polyspecific TCR

To determine the molecular basis for the polyspecificity of these TCR clonotypes, we developed a library-on-library scheme that would enable us to map T cell epitope libraries against a T cell hybridoma library efficiently (**Fig. 4a**). We generated genomic libraries in *E. coli* from three of the strains that stimulated all the Firmicute-reactive hybridomas: *Clostridium bolteae*, *Tyzzerella nexilis*, and *Subdoligranulum variabile*. We organized each genomic library into 480 pools of 30 clones each and screened the pools against 13 co-cultured hybridomas. We found a rare positive clone pool from each of the three libraries; a subsequent deconvolution step yielded individual stimulatory clones from *C. bolteae*, *T. nexilis*, and *S variabile*. Strikingly, these three clones are related in primary amino acid sequence (**Fig. 4b**): each of them contains a region from the C-terminal domain of SBP that is predicted to function as part of an ABC transport system for monosaccharide utilization (**Fig. 4c**). The SBP of gram-positive bacteria is an extracellular lipoprotein that is anchored in the outer leaflet of the plasma membrane^36,37^. Two computational tools predicted that the SBP from *T. nexilis* has a lipoprotein signal peptide (**Extended Data Fig. 5a, b**). The predicted extracellular localization of the SBP may contribute to its strong antigenicity by making the antigen more accessible to immune cells. Of note, among the strains in hCom1d and hCom2d, there is a near-perfect correspondence between the presence of the SBP in the genome and stimulation of the hybridomas in question; the only exception is the actinobacterium *Collinsella stercoris* DSM 13279 (**Fig. 4d** and **Extended Data Fig. 5c**). Although they are all Firmicutes (except *Collinsella stercoris* DSM 13279), they do not derive from a monophyletic clade (**Fig. 4e**), so the trait(s) they share appear to be shaped by gene acquisition or loss.

**Figure 4:**
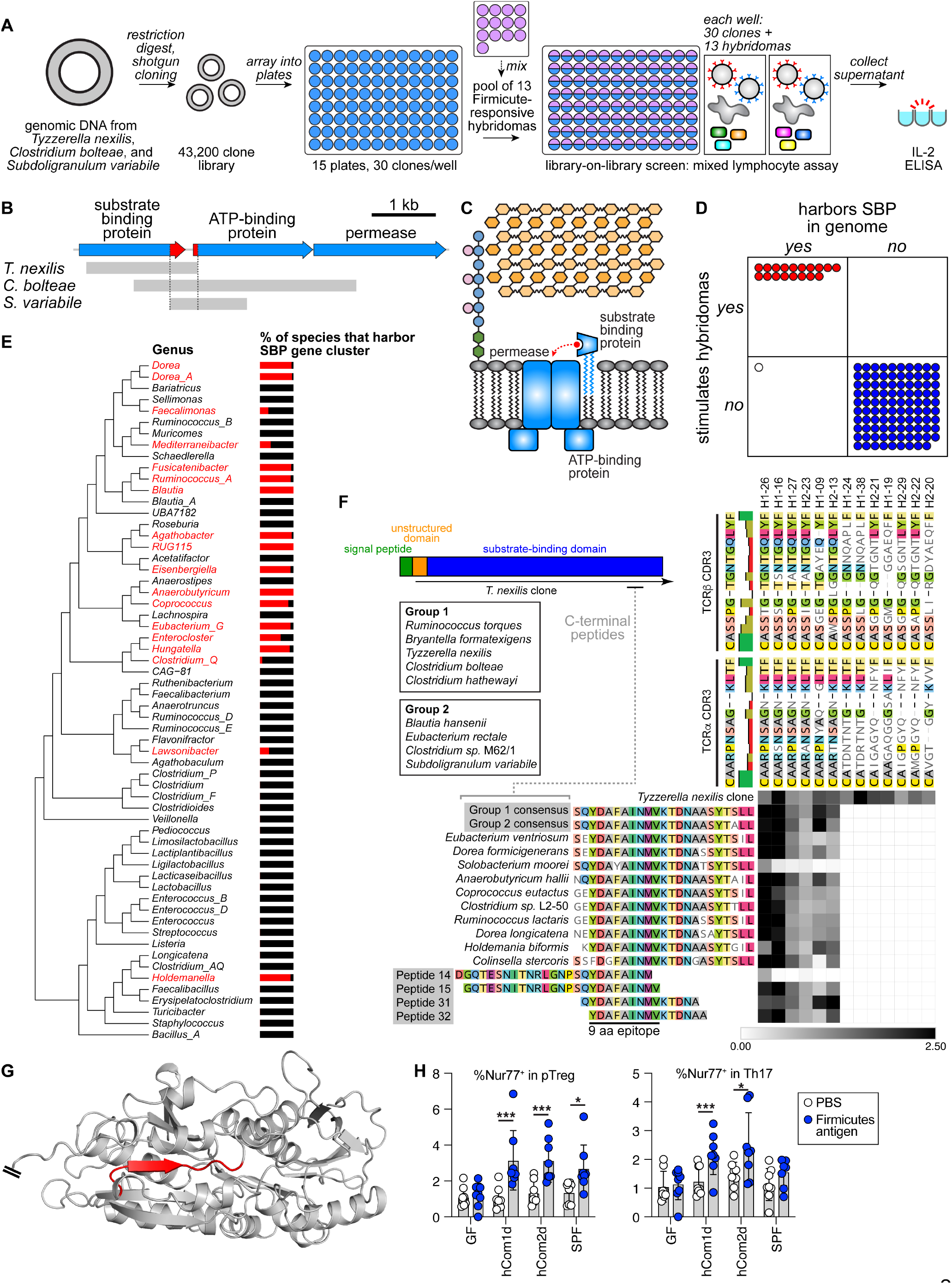
Discovery of a conserved Firmicutes antigen. (**A**) Schematic of the antigen identification experiment. Genomic DNA was extracted from *Tyzzerella nexilis, Clostridium bolteae,* and *Subdoligranulum variabile,* partially digested by a restriction enzyme, cloned into the pGEX-4T1 expression vector, and electroporated to *E. coli* to generate three genomic libraries. Each of three genomic libraries was cultured in plates as 480 pools of 30 clones each (14,400 *E. coli* clones per library, 43,200 in total). Each library was screened against a mixture of 13 Firmicutes-reactive hybridomas (**Fig. 3e**). To monitor TCR simulation, we measured IL-2 production by ELISA. (**B**) One positive clone was identified from each of the three genomic libraries; the insert in each clone was analyzed by Sanger sequencing. The genomic fragments in each clone overlap around the C-terminal domain of a SBP that is predicted to function as part of an ABC transport system for monosaccharide utilization. (**C**) Schematic of the ABC transport system and SBP. The SBP is predicted to be a lipoprotein that is anchored in the outer leaflet of the plasma membrane. (**D**) BLAST-based genomic analysis of SBP distribution in hCom strains. The presence of an SBP homologous to those found in *T. nexilis, C. bolteae,* and *S. variabile* is almost perfectly correlated to the ability of a strain to stimulate a subset of 13 hybridomas as shown in **Fig. 3E**. (**E**) A phylogenetic tree showing the distribution of the SBP gene cluster among host-associated Firmicutes. Genera colored red include species that harbor the SBP cluster; the percentage of species that harbor the cluster is indicated by the red portion of the bar to the right of each genus. (**F**) Identification of Firmicutes antigen epitope 1 (FA1). C-terminal SBP peptides from 19 stimulatory strains from hCom1d and hCom2a were synthesized and co-cultured with the T cell hybridomas. The truncated peptides 14, 15, 31 and 32 were also tested in order to define the minimal epitope. 6 of the 13 TCRs were responsive to the synthetic peptides. The remaining 7 were stimulated only by an *E. coli* clone harboring nearly all of the SBP sequence from *T. nexilis,* indicating the presence of an additional epitope(s). There is a strong correlation between the reactivity of TCRs and the sequences of TCR CD3 regions. Consistent with the data in **Fig. 3E**, the peptides from *Solobacterium moorei* did not stimulate TCRs. (**G**) Predicted crystal structure of the SBP from AlphaFold2. The FA1 epitope, shown in red, lies within the central beta-sheet of the membrane-proximal domain. (**H**) Induction of SBP-specific T cells *in vivo.* Germ-free C57BL/6 mice were colonized with hCom1d or hCom2d. After two weeks, intestinal T cells were isolated and cocultured with FA1 and dendritic cells. Nur77 expression in T cell subsets was analyzed by FACS to monitor TCR stimulation. In hCom1d and hCom2d-colonized mice, Th17 and Treg cells showed an antigen-specific response to SBP. Notably SBP-specific T cells were also found in SPF mice, which are colonized by an entirely murine microbiota; suggest that FA1 is broadly conserved among Firmicutes that colonize the intestine.

To narrow down the region within the C-terminus that harbors the MHC II epitope, we synthesized 42 peptides from the *T. nexilis* SBP that tile the 53 amino acid region of overlap among the three clones from the screen. We co-cultured each candidate peptide with a pool of 13 mixed TCR hybridomas (**Extended Data Fig. 6**). Truncated peptides containing the 9-mer YDAFAINMV fully stimulated the mixed hybridomas, whereas peptides that lack any of these residues are inactive or only weakly stimulatory.

This epitope is identical among the Firmicutes that were found to stimulate this subset of hybridomas but the surrounding sequence is not. We tested twelve 22-23-mer peptides surrounding this epitope, representing the SBP sub-sequences from all the stimulatory Firmicutes, against each of the 13 hybridomas (**Fig. 4f**). This experiment yielded two findings: First, we confirm that the epitope is YDAFAINMV (hereafter, Firmicutes antigen epitope 1, FA1). In support of this assignment, *Solobacterium moorei* has an F to Y mutation in this region; as a result, the *S. moorei* peptide only weakly stimulates the corresponding T cell hybridomas. An AlphaFold2-based prediction of the SBP structure suggests that this epitope lies within the central beta-sheet of the membrane-proximal domain (**Fig. 4g**), where amino acid sequences are more conserved than in variable loop regions, providing one possible explanation for the degree of its conservation.

Second, YDAFAINMV is the target of only 6 of the 13 hybridomas. Notably, an *E. coli* clone harboring the genomic region from *T. nexilis—*which covers most of the amino acid sequence of the SBP— stimulates all 13 hybridomas. These data suggest that the remaining 7 hybridomas are specific for different epitope(s) from the same protein.

Finally, we sought to determine the prevalence of FA1-specific T cells in the gut repertoire. We isolated T cells from hCom1d- and hCom2d-colonized mice, SPF mice, and germ-free mice as a control (**Fig. 4h**). As expected, FA1 restimulates pTreg and Th17 cells from hCom1d and hCom2d-colonized mice.

Notably, FA1 also stimulates pTreg cells from SPF mice. These mice are colonized by an entirely murine microbiota; they have little overlap in microbial composition with hCom1d or hCom2d (which are human isolates). Our results suggest that the broad conservation of FA1 among a subset of Firmicutes makes FA1-specific T cells a substantial portion of the gut repertoire, even in mice who do not harbor the bacterial strains from which FA1 was discovered.

## DISCUSSION

The system we introduce here has three features that make it powerful for studying immune modulation by the gut microbiome. First, the gut community is defined, enabling the interrogation of individual strains to quantify their contribution to a community-level phenotype. In this way, we identified a variety of strains capable of restimulating Treg and Th17 cells (**Fig. 2**); these are a promising starting point for identifying new antigen-specific T cell inducers that function robustly in the setting of a complex community. Although we focused here on mapping strains to the T cell repertoire, our system should be equally useful in identifying causative strains for an immune modulatory phenotype, e.g., by strain dropout from (or addition to) a defined community.

Second, the use of scRNA-seq and scTCR-seq, in conjunction with T cell hybridoma construction, enabled us to construct a map of strains to TCR clonotypes. Two factors were important: the expansion of TCR clonotypes in colonized mice was a useful predictor of microbiome specificity, and a protocol for the rapid construction of >90 hybridomas in parallel—by enrichment rather than cell sorting—enabled us to test TCRs at scale. The combination of scRNAseq and scTCRseq has been used in prior work^29,38^, but studies have been limited to the comparison of TCR sequences and have not yet identified the antigens they recognize. Our approach provides a method for mapping dozens of TCRs to antigens in parallel.

Strikingly, among 31 TCRs reactive to bacterial strains, 30 reacted to multiple strains. This pattern deviates from the current model, in which T cells are specific for an individual colonist^1–4^. Instead, it suggests that in the physiological setting of a complex microbial community, expanded TCR clonotypes play ‘zone defense’—instead of recognizing a single colonist they mediate the response to a group of colonists that share a conserved epitope. Most polyreactive TCRs targeted strains of Firmicutes or Bacteroidetes, as observed recently in a different setting^32^ and a distinction from previous work reporting poly-reactive immunoglobulin A (IgA) that recognizes strains of Proteobacteria^39^. Future experiments are needed to find dietary antigens for the four TCRs that were specific to chow and fecal pellets from germ-free mice.

Third, in cloning the epitope recognized by the Firmicutes-specific T cell hybridomas, it was essential that our pool of possible TCR epitopes was completely defined. Each strain in the community is sequenced and arrayed as a pure culture, enabling the construction of genomic libraries from strains of interest. Screening pools of genomic clones against pooled hybridomas made it possible to test 43,200 clones against 13 hybridomas (561,600 combinations) in just 1,440 ELISA assays (arrayed across 15 plates), leading to the identification of three clones—one from each genomic library—which encoded overlapping fragments of orthologous SBPs from an ABC transport system. The SBP is extracellular, widely conserved among Firmicutes, and presumably highly expressed, features that are likely to be found in other epitopes recognized by polyspecific T cells that patrol the gut microbiome^32^.

We hypothesize that a T cell clone specific for the conserved SBP epitope, FA1, may expand quickly because it encounters SBP-expressing Firmicutes frequently at barrier surfaces along the intestine. Notably, the Firmicutes that stimulated ‘polyspecific’ clonotypes are not the most abundant strains in the community. We speculate that they might have privileged access to a barrier surface, or they produce a metabolite that facilitates immune modulation.

If the polyspecificity of microbiome-directed TCRs is borne out by subsequent studies, it will have important implications for the logic of immune surveillance against the gut microbiota. A typical gut community consists of a massive number of strains and epitopes; mounting a prophylactic defense against all of them, using a limited set of lymphocyte clonotypes, is an imposing challenge for the host. Our data suggest that the T cell component of the immune response addresses this challenge by playing ‘zone defense’: clonotypes that recognize broadly conserved antigens undergo positive selection, providing single-clonotype defense against a swath of strains. This model echoes some of the characteristics of broadly neutralizing antibodies against HIV and SARS-Co-V2^40,41^, which are specific at the level of molecular recognition but provide broad protection against pathogens because their target is widely conserved across variants.

Our work has three important limitations. First, we analyzed gnotobiotic mice after two weeks of community colonization, a timepoint when adaptive immune responses are at their peak, to maximize the likelihood of finding microbiome-specific T cells. To characterize physiological immune development, a different time point (e.g., colonize at weaning and sample 2-3 months later) would be useful to assess in the future.

Second, although FA1 is conserved across 18 Firmicutes species from hCom1d and hCom2d, it is unclear whether all 18 strains contribute to FA1 -specific T cell responses in vivo or a subset are dominant. This question could be addressed in future work by constructing dropout communities in which a subset of the 18 strains are missing and assessing the number and subtype of FA1-specific T cells. A similar approach could be used to investigate whether the mixture of Tregs and T effector cells observed among FA1-specific T cells arises from individual strain(s) that induce both phenotypes vs. multiple strains that induce a single T cell subtype.

Third, the function of FA1-specific T cells is not yet known. Given their abundance, they have the potential to play an important role in immune response to commensals. Moreover, they are a mixture of Treg and Th17 cells, raising the possibility that the balance within the FA1-specific pool might be relevant to inflammatory and autoimmune disease^5^.

An epitope that is the focus of a concerted T cell response against the microbiome creates new therapeutic opportunities. By rationally altering the composition of the community, it may be possible to influence the balance of Treg and effector T cells in the FA1-specific pool. Alternatively, key FA1-harboring strains could be engineered to express non-native antigens, redirecting this pool against a target of therapeutic interest^42^. Finally, it might be possible to engineer FA1-specific T cells directly using new approaches to target and transduce T cells based on their TCR specificity^43,44^. These approaches could open the door to new therapeutic communities for inflammatory bowel disease, cancer, infectious disease and autoimmune disease.

## MATERIALS AND METHODS

### Bacterial strains and synthetic community construction

hCom1d and hCom2d were prepared from individual strain stocks as described previously^26^. For germ-free mouse experiments, strains were revived from frozen stocks and cultured for 48 h anaerobically in an atmosphere consisting of 10% CO2, 5% H2, and 85% N2. Strains were propagated in sterile 2.2-mL 96-well deep well plates in their respective growth medium (**Supplementary Table 1**): Mega Medium^26^ supplemented with 400 μM vitamin K2,or Chopped Meat Medium supplemented with Mega Medium carbohydrate mix^26^ and 400 μM vitamin K2. The optical density at 600 nm (OD600) of each revived strain was measured. We pooled appropriate volumes of each culture corresponding to 2 ml at OD_600_=1.3, centrifuged for 5 min at 5000 x *g,* and resuspended the pellet in 2 m. of 20% glycerol. For each inoculum preparation cycle, a small number of strains typically did not reach OD_600_ ~1.3. For these strains, the entire 4 ml culture volume was added to the pooled strain mixture. Following pooling and preparation, 1.2 ml of the synthetic community was aliquoted into 2 ml cryovials and stored at −80 °C. To colonize C57BL/6 germ-free mice, a single cryovial was thawed and 200 μl was administered by oral gavage into 8-12-week-old mice. 2 weeks after colonization, mice were sacrificed for analysis as detailed below.

To prepare heat-killed bacterial strains for in vitro co-culture experiments, individual strains were revived from frozen stocks under the anaerobic conditions described above. After measuring the OD600, each culture was centrifuged for 5 min at 5000 x *g* and resuspended into a volume of PBS required to normalize OD600 to 1.0. Normalized cultures were heat-treated in a water bath at 70 °C for 30 min and then stored at −30 °C.

Metagenomic sequencing and sequence analysis were performed as described previously^26^. In brief, fecal pellets were collected from mice into sterile tubes. Genomic DNA was extracted using the DNeasy PowerSoil HTP kit (Qiagen) and quantified in 384-well format using the Quant-iT PicoGreen dsDNA Assay Kit (Thermofisher). Sequencing libraries were generated in 384-well format using a custom low-volume protocol based on the Nextera XT process (Illumina). Sequencing reads were generated using a NovaSeq S4 flow cell or a NextSeq High Output kit, in 23×150 bp configuration. 5-10 million paired-end reads were targeted for isolates and 20-30 million paired-end reads for communities. NinjaMap^26^ was used to calculate relative abundance of each strain.

### Isolating immune cells from the intestinal lamina propria

Mice were euthanized by cervical dislocation. The small and large intestine were collected by dissection and the mesenteric lymph nodes, Peyer’s patches, and cecum were removed. Intestinal tissue was shaken at 225 rpm for 40 min at 37 °C in DMEM (Gibco) containing 5 mM EDTA and 1 mM DTT. Tissues were washed with DMEM, manually shaken to detach the epithelial layer and cut into pieces. Tissue fragments were then digested using a mouse Lamina Propria Dissociation Kit (Miltenyi Biotec, #130-097-410) and a gentleMACS dissociator (Miltenyi Biotec). After digestion, debris was removed using a 40% and 80% Percol gradient purification (GE Healthcare).

### Induction of splenic DCs by injecting B16-FLT3L

8-12-week-old CD45.1^+^ C57BL/6 mice were purchased from the Jackson Laboratory (Stock No: 002014). 5.0 x 10^6^ B16-FLT3L cells^45^ were administered to mice by intraperitoneal injection to expand FLT3L-dependent DCs. Mice were sacrificed 12-14 d after injection. Spleens were excised and digested using a spleen dissociation kit (mouse) (Miltenyi Biotec, #130-095-926) and a gentleMACS dissociator (Miltenyi Biotec). Red blood cells were lysed and CD11c^+^ dendritic cells were enriched using CD11c MicroBeads UltraPure (mouse) (Miltenyi Biotec, #130-125-835).

### Co-culture of primary intestinal immune cells with splenic DCs and antigens

5.0 x 10^5^ CD45.1^+^ FLT3L-induced DCs were co-cultured in DMEM + 10% FBS with one of the following stimulants for 30 min at 37 °C in a 5% CO2 incubator: 5 μl PBS, 5 μl of a synthetic antigen peptide (Genscript, 0.2 mg/ml in PBS), eBioscience Cell Stimulation Cocktail (Fisher Scientific 00-4975-93) or 5 μl of heat-killed bacteria in PBS, prepared as described above. After 30 min, 5.0 x 10^4^ freshly isolated cells from the intestinal lamina propria were added and co-cultured for another 4 h before staining and fixation. To screen T cell responses to bacterial strains from hCom1d, we pooled T cells from 10-15 mice in each experiment.

### Cell fixation, staining and flow cytometry

Single cell suspensions were stained with Fixable Viability Dye eFluor 780 (eBioscience, 65-0865-18) before fixation to detect dead cells. Cells were then fixed for 30 min at room temperature using the fixation/permeabilization buffer supplied with the eBioscience Foxp3/Transcription Factor Staining buffer set (ThermoFisher Scientific, 00-5523-00).

After fixation, cells were stained with combinations of the following primary antibodies in permeabilization buffer for 30 min at room temperature: Helios (22F6, BioLegend), T-bet (4B10, BioLegend), Nur77-PE (12.14, eBioscience), Foxp3 (FJK-16s, eBioscience), Nur77-purified (11C1052, LSBio), CD45.1 (A20, eBioscience), CD4 (RM4-5, BioLegend), CD3e (145-2C11, BioLegend), FR4 (12A5, BD), IL17A (eBio17B7, eBioscience), RORγt (B2D, eBioscience, or Q31-378, BD Bioscience), IFNγ (XMG1.2, BioLegend), PD-1 (29F.1A12, BioLegend) and IL10 (JES3-9D7, BioLegend). Cells stained with the Nur77-purified antibody (11C1052, LSBio) were further stained with a secondary APC Goat anti-Rabbit IgG (Polycolonal, eBioscience). All staining was performed in the presence of purified anti-mouse CD16/32 (clone 93) and 10% fetal bovine serum to block nonspecific binding. Cells were analyzed using an LSR II flow cytometer (BD Biosciences) and data were processed using FlowJo (TreeStar).

### scRNA-seq and scTCR-seq analysis

We colonized 8-10-week-old C57BL/6 germ-free mice with hCom1d or hCom2d. We pooled three gender-matched mice per condition, isolated lymphocytes from the lamina propria of the small and large intestine, and enriched the live CD3^+^ T cell fraction by cell sorting (FACS Aria II, BD Biosciences). Sorted cells were re-suspended in PBS with 0.05% BSA and ~1 × 10^4^ cells were loaded onto the Chromium controller (10x Genomics). Chromium Single Cell 5’ reagents were used for library preparation according to the manufacturer’s protocol. The libraries were sequenced on an Illumina HiSeq 4000. Sequencing data were aligned to the reference mouse genome mm10 with Cell Ranger (10x Genomics). The data were processed using the R packages Seurat version 3.1.5^46^ and scRepertoire version 1.0.0^47^.

For scRNA-seq analysis, we excluded cells with fewer than 500 and more than 4500 detected genes. We further eliminated cells with >25% mitochondrial genes. Data were clustered using the FindClusters function of Seurat. Visualization of the clusters on a 2D map was performed with uniform manifold approximation and projection (UMAP) (RunUMAP function of Seurat, dims = 1:20). To assign cell clusters to T cell subsets, gene expression levels in clusters were visualized by dot plots, violin plots and feature plots using Seurat. Differentially expressed genes of each T cell subset between colonization conditions were identified using the FindMarkers function of Seurat.

scRepertoire was used to merge scTCR-seq data with the Seurat object of scRNA-seq data. The expansion of TCR clonotypes was visualized on UMPA by the DimPlot function of Seurat (group.by = cloneType). To characterize T cell phenotypes of TCR clonotypes, a metadata file was generated from the Seurat object and analyzed and quantified using Microsoft Excel.

### Generating TCR hybridomas

We generated the retroviral vector pMSCV-mCD4-PIG TCR-OTII to transduce NFAT-GFP hybridoma cells with synthetic TCR constructs. pMSCV-PIG (#21654) was purchased from Addgene, subjected to restriction digest by BglII, and purified. To generate pMSCV-PIG TCR-OTII, a gBlock encoding the OTII TCR was ordered from IDT and ligated into BglII-digested pMSCV PIG using HiFi DNA assembly master mix (E2621S, New England Biolabs). pMSCV-PIG TCR-OTII was digested with BstXI and SalI and then ligated with a gBlock encoding the mammalian CD4 gene to make pMSCV-mCD4-PIG TCR-OTII. Plasmid and gBlock sequences are listed in **Supplementary Table 3**.

pMSCV-mCD4-PIG TCR-OTII, our backbone vector for TCR expression, was provided to Twist Bioscience. Sequences of the TCR a and β genes we selected (92 TCRs in total) were obtained from the JSON files generated by scTCR-seq. A cassette that consists of the TCR a and β genes separated by a self-cleaving P2A peptide was synthesized (Twist Bioscience) and used to replace the OTII TCR gene in pMSCV-mCD4-PIG.

The Platinum-E (Plat-E) retroviral packaging cell line (RV-101, Cell Biolabs) was used for viral packaging of pMSCV-mCD4-PIG TCR vectors. Plat-E cells were cultured in a 96-well plate to 60-80% confluence and transfected with pMSCV-mCD4-PIG TCR vectors as a DNA-lipid complex using Lipofectamine 3000 (L3000001, ThermoFisher Scientific) following the manufacturer’s instructions. 24 h after transfection, culture supernatant containing viral particles was harvested.

2.5 × 10^4^ NFAT-GFP cells were centrifuged, resuspended in 200 μl of the virus-containing supernatant supplemented with 10 μg/ml protamine sulfate (MP Biomedicals), and plated in u-bottom 96-well plates. NFAT-GFP cells were spin-transduced by centrifugation at 1000 x *g* at 32 °C for 120 min, then pipetted up and down and cultured overnight at 37 °C in a 5% CO2 incubator. Cells were then transferred into a flask and incubated for 1-2 weeks with 10 ml of culture medium (DMEM with 10% FCS, Pen/Strep and 2 mM L-glutamine) under puromycin selection (1 μg/ml). After selection and propagation, TCR expression by NFAT-GFP cells was confirmed by flow cytometry.

### TCR hybridoma co-culture experiment

2.0 × 10^4^ TCR-transduced NFAT-GFP cells were co-cultured in a 96 well plate with 1.0 × 10^5^ FLT3L-induced DCs and one of the following stimuli at 37 °C in a 5% CO2 incubator: 5 μl PBS, 5 μl synthetic antigen peptide (Genscript, 0.2 mg/ml in PBS), or 5 μl heat-killed bacteria in PBS. After 24 h of co-culture, cells were centrifuged and culture supernatant was collected. TCR stimulation was evaluated by measuring IL-2 concentration in culture supernatant using an IL2 ELISA kit (#423001, #423501, #431001 and #421101, BioLegend).

### Antigen discovery using *E. coli* shotgun genomic libraries

*E. coli* shotgun genomic libraries were generated as described^1,33,48^ with minor modifications. Bacterial genomic DNA was purified from liquid cultures of *Tyzzerella nexilis, Clostridium bolteae,* and *Subdoligranulum variabile* using a QIAamp PowerFecal DNA Kit (Qiagen). 0.1 mg of genomic DNA per strain was partially digested by Sau3AI. For partial digestion, Sau3AI was serially diluted and aliquots of genomic DNA were added. After 10 min of digestion, Sau3AI was heat-inactivated. All the reactions from the serially dilutions of Sau3AI were mixed after digestion and subjected to gel electrophoresis. DNA fragments between 500-5,000 bp were gel purified. The pGEX-4T1 expression vector (GE28-9545-49,Sigma-Aldrich) was digested with BamHI and dephosphorylated using the Quick Calf Intestinal Alkaline Phosphatase kit (M0525L, New England Biolabs). CIP was heat-inactivated and the linearized pGEX-4T1 vector was gel-purified.

Sau3AI-digested bacterial genomic DNA and the linearized, dephosphorylated pGEX-4T1 backbone were ligated using the T4 DNA Ligase kit (M0202M, New England Biolabs). Ligation products were transformed into ElectroMAX DH10B competent cells (18290015, Thermo Fisher Scientific) by electroporation. DH10B cells were plated on LB-agar + 100 μg/ml carbenicillin. CFUs were counted and several colonies were picked for Sanger sequencing to verify the presence of a bacterial genomic fragment. DH10B cells were then adjusted to 30 CFUs per 100 μl of LB medium containing 10% glycerol (v/v) and 100 μg/ml carbenicillin and cultured in 96-well plates overnight. 5 plates were prepared per bacterial strain to generate shotgun genomic libraries. 7 μl of DH10B culture per well was sub-cultured in 200 μl of fresh LB-carbenicillin medium in 96 well plates to an OD600 of 0.5-0.6; the remained of the DH10B culture was stored at −20 °C. IPTG (Sigma-Aldrich) was added to a final concentration of 0.5 mM and DH10B cells were incubate for 16 h at 18 °C to induce the expression of inserted bacterial genes. DH10B cells were centrifuged, washed with PBS, heat-killed by incubating at 75 °C for 1 h and stored at −20 °C until use.

For co-culture experiments, 13 Firmicutes-reactive NFAT-GFP hybridomas were mixed. 2.0 x 10^5^ mixed NFAT-GFP cells were co-cultured in 150 μl medium (DMEM with 10% FCS, Pen/Strep) in a 96-well plate with 2.0 × 10^5^ FLT3L-induced DCs and 5 μl of heat-killed DH10B cells in PBS. After 24 h of co-culture, cells were centrifuged and culture supernatant was collected to measure IL-2 concentration as a readout of TCR stimulation.

### Protein Structure Prediction by AlphaFold2 with MMseqs2

To predict the 3D structure of the substrate binding protein, we used ColabFold: AlphaFold2 protein structure^49^ and complex^50^ prediction using multiple sequence alignments generated through MMseqs2^51^. Predicted structures were visualized by PyMOL v2.5 (Schrödinger, Inc).

### Phylogenetic distribution of the SBP gene cluster

Metadata was obtained for all genomes in the UHGG v2 database as of March 25, 2022 was obtained from their FTP site (https://ftp.ebi.ac.uk/pub/databases/metagenomics/mgnify_genomes/human-gut/v2.0/genomes-all_metadata.tsv). This list was filtered such that only the isolate genomes that belonged to the phylum Firmicutes remained—2,651 genomes across 237 genera remained after filtering. The list was filtered further to keep only those genera that contained at least 10 isolate genomes. Our final dataset contained 2,174 genomes from 60 Firmicutes genera. These 2,174 genomes and their genomic feature files were downloaded from the FTP site. All genomic features were extracted from the genomes using the “getfasta” module from the bedtools toolkit. A representative genome from each genus was picked to generate a tree using GTDBtk classify workflow using the release202 version of the database.

The three proteins from the SBP operon were extracted from the genome sequence of *Clostridium nexile* DSM 1787 and a protein blast database was created using these sequences. BlastX was run with all the genes from the 2,174 genomes as query and the proteins from the operon as the database (non-default settings: -dbsize 1000000 -num_alignments 5 -outfmt ‘6 std qlen slen qcovs ppos’). Another BlastX search was run with the genes from the Firmicutes in hCom1d and hCom2d using the same database and blast settings. The blastx results were filtered such that only those hits were considered that had a bit score >= 560 and percent identity >= 60%. Finally, of the 386 UHGG genomes that contained a hit that passed our thresholds, 366 genomes across 18 genera contained proteins homologous to the proteins in the SBP operon. A custom R script (github: decorate_tree.Rmd) was used to integrate the UHGG blast results and the genera tree created from GTDBtk.

### Statistics

All statistical analyses was performed with Graphpad Prism9 (GraphPad Software, La Jolla, CA). P values < 0.05 were considered statistically significant.

## ACKNOWLEDGMENTS

We are deeply indebted to Djenet Bousbaine and other members of the Fischbach Group for helpful suggestions and comments on the manuscript. We thank Hiroshi Takayanagi, Shinichiro Sawa, Ryunosuke Muro and members of their labs for useful discussions. We are grateful to the Stanford Shared FACS Facility for an access to flow cytometry and the Stanford Genomics Facility for constructing and sequencing libraries for scRNA-seq and scTCR-seq experiments.

This work was supported by the Stanford Microbiome Therapies Initiative, the Human Frontier Science Program LT000493/2018-L (K.N.), a Fellowship from the Astellas Foundation for Research on Metabolic Disorders (K.N.), a research grant from Kanae Fundation for the Promotion of Medical Science (K.N.), an HHMI-Simons Faculty Scholar Award (M.A.F.); the Leona M. and Harry B. Helmsley Charitable Trust (M.A.F.); NIH grant DK110174 (M.A.F.); the Chan Zuckerberg Biohub (M.A.F.); Stand Up to Cancer (M.A.F.); and the MAC3 Impact Philanthropies (M.A.F.).

## AUTHOR CONTRIBUTIONS

K.N., and M.A.F. conceived and designed the experiments. K.N., A.Z., K.A., A.W., S.J., X.M., A.G.C., M.W., S.H., A.D., and P.M. performed the experiments. K.N. and M.A.F. analyzed data and wrote the manuscript. All authors discussed the results and commented on the manuscript.

